# Disabling *de novo* DNA methylation in embryonic stem cells allows an illegitimate fate trajectory

**DOI:** 10.1101/2020.12.20.423404

**Authors:** Masaki Kinoshita, Meng Amy Li, Michael Barber, William Mansfield, Sabine Dietmann, Austin Smith

## Abstract

Genome remethylation is essential for mammalian development but specific reasons are unclear. Here we examined embryonic stem (ES) cell fate in the absence of *de novo* DNA methyltransferases. We observed that ES cells deficient for both *Dnmt3a* and *Dnmt3b* are rapidly eliminated from chimaeras. On further investigation we found that in vivo and in vitro the formative pluripotency transition is derailed towards production of trophoblast. This aberrant trajectory is associated with failure to suppress activation of *Ascl2. Ascl2* encodes a bHLH transcription factor expressed in placenta. Misexpression of *Ascl2* in ES cells provokes transdifferentiation to trophoblast-like cells. Conversely, *Ascl2* deletion rescues formative transition of *Dnmt3a/b* mutants and improves contribution to chimaeric epiblast. Thus, *de novo* DNA methylation safeguards against ectopic activation of *Ascl2*. However, *Dnmt3a/b*-deficient cells remain defective in ongoing embryogenesis. We surmise that multiple developmental transitions may be secured by DNA methylation silencing potentially disruptive genes.

**SIGNIFICANCE STATEMENT:** Mammalian DNA is widely modified by methylation of cytosine residues. This modification is added to DNA during early development. If methylation is prevented, the embryo dies by mid-gestation with multiple abnormalities. In this study we found that stem cells lacking the DNA methylation enzymes do not differentiate efficiently into cell types of the embryo and are diverted into producing placental cells. This switch in cell fate is driven by a transcription factor, Ascl2, which should only be produced in placenta. In the absence of DNA methylation, the *Ascl2* gene is mis-expressed. Removing Ascl2 redirects embryonic fate but not full differentiation potential, suggesting that methylation acts at multiple developmental transitions to restrict activation of disruptive genes.

## INTRODUCTION

The mammalian genome is characterised by widespread methylation of cytosine residues. After fertilisation, however, both maternal and paternal genomes undergo extensive demethylation, reaching a low point in the blastocyst (1-4). The embryo genome is then remethylated by the activity of de novo DNA methylation enzymes (5). Mouse embryonic stem (ES) cells exhibit global hypomethylation, similar to the in vivo blastocyst profile (6-8). Methylation is gained during the formative pluripotency transition to lineage competence, recapitulating early post-implantation development in vivo (9, 10).

Mammals have three DNA methyltransferases (DNMTs). Dnmt1 maintains methylation during DNA replication, while Dnmt3a and Dnmt3b are responsible for de novo methylation. DNA methylation is not required for general cell viability and, with the exception of imprint control regions, largely occurs at seemingly non-functional regions of the genome (11). Nonetheless, knockout of *Dnmt1* in mice results in embryonic lethality around E9.5 (12). *Dnmt3a* mutants die during puberty, but *Dnmt3b* mutant embryos fail from E9.5 onwards, exhibiting multiple abnormalities (13). When both de novo DNMTs are inactivated, development is disrupted by E8.5 with defective somitogenesis and abnormal morphogenesis. Seemingly normal progress to late gastrulation suggests that remethylation in the early post-implantation embryo is not critical for epiblast transition or germ layer formation. However, the reason(s) for the subsequent catastrophic failure is unclear. It has also been found that ES cells doubly deficient for *Dnmt3a* and *Dnmt3b* show a progressive genome-wide reduction in DNA methylation and loss of ability to form teratomas after long-term culture (14).

We previously showed that depletion of Dnmt3a and Dnmt3b delays timely exit from naïve pluripotency in vitro (15). Here we investigate the functional consequences of lack of de novo methylation in ES cells for pluripotency progression and lineage potential at the cellular level.

## RESULTS

### Chimaera colonisation and lineage potential of Dnmt3a/3b deficient ES cells

We examined the ability of *Dnmt3a* and *Dnmt3b* double knockout (Dnmt3dKO) ES cells (15) to contribute to chimaeric embryos. Compound *Dnmt3a/3b* mutant embryos are reportedly normal until somitogenesis (13). However, after blastocyst injection of Dnmt3dKO cells genetically labelled with constitutive mKO2, we found very sparse contributions in pre-somite stage embryos at E7.5 (Fig. 1A). Even at E6.5 contributions were reduced compared with typical ES cell chimaeras (Fig. 1B). Furthermore, some mutant donor cells were located in the extraembryonic ectoderm, a rare occurrence with wild type ES cells (16).

**Figure 1.**
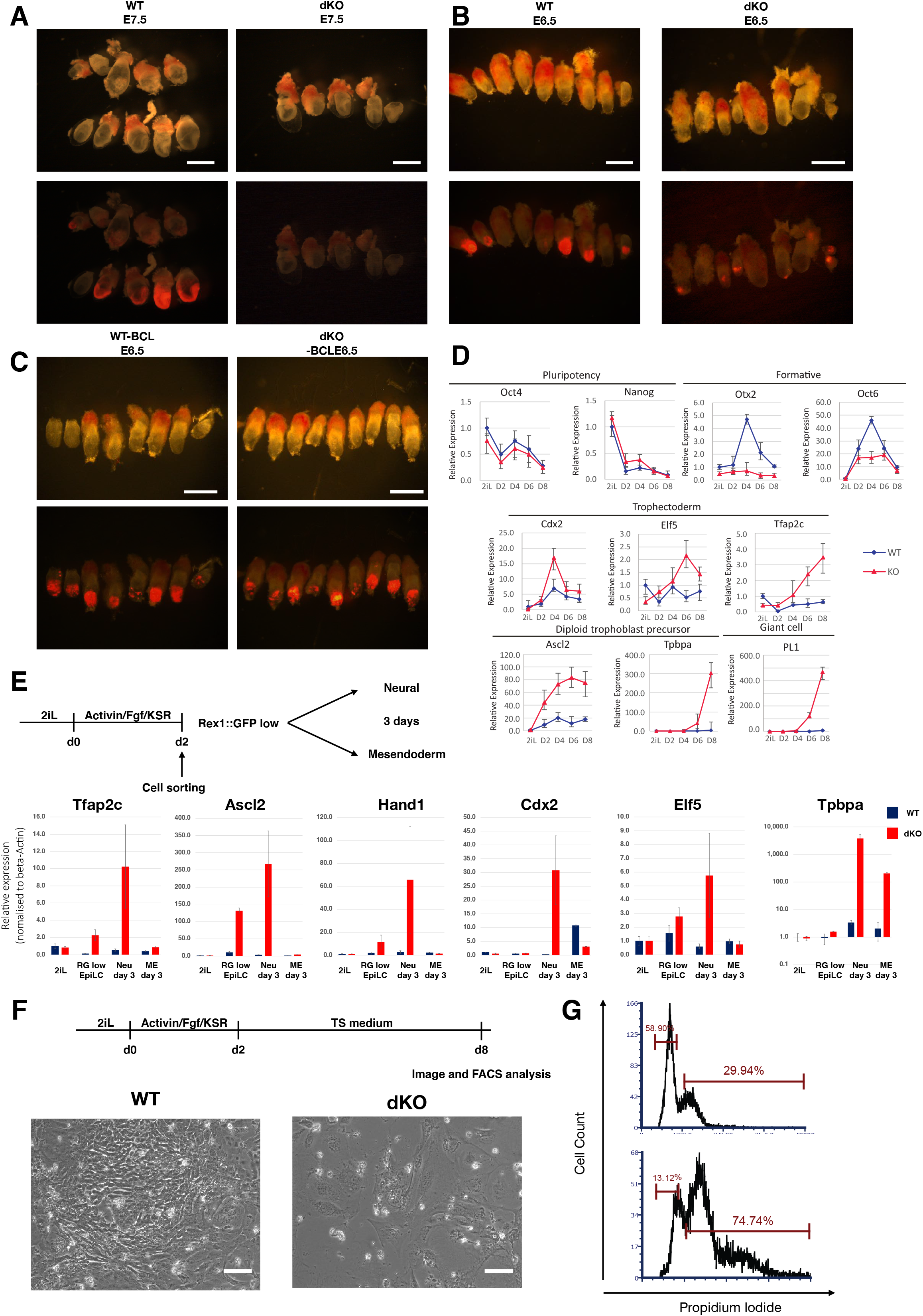
*Dnmt3a/b*-deficiency compromises chimaera contribution and somatic lineage commitment. (A-C) Chimaeric embryos obtained with parental, Dnmt3dKO (A, B), and hBCL2 expressing (C) mKO2 reporter ES cells. 30% opacity bright field images overlaid onto fluorescent images. Scale bars, 1mm (A), 500µm (B-C). (D) qRT-PCR analysis of marker during culture in trophoblast medium (20). (E) qRT-PCR analysis of undifferentiated ES cells, Rex1::GFP low sorted AFK cells, and further differentiated cells. (F) Cell morphology of WT and Dnmt3dKO cells in TS cell medium for 8 days. Scale bars, 50µm. (G) DNA content quantified by propidium iodide (PI) staining at day 8 in TS cell medium. qRT-PCR data were normalized to beta-Actin. Error bars are S.D. from technical duplicates.

To improve survival of Dnmt3dKO ES cells in chimaeras we introduced a constitutive *BCL2* transgene (17). In E6.5 embryos we observed higher contributions to epiblast, comparable to wildtype ES cells (Fig. 1C). However, we also saw donor cells in extraembryonic regions. We confirmed localization in extraembryonic ectoderm and ectoplacental cone in 5 out of 6 chimaeras examined by confocal microscopy (Fig S1A). We inspected blastocyst stage chimaeras to test whether *Dnmt3a/b* deficiency or *BCL2* expression enabled primary trophectoderm colonization. However, donor cells were correctly localized to the inner cell mass and did not contribute to trophectoderm (Fig. S1B). Thus, the presence of Dnmt3dKO cells in post-implantation trophoblast likely arises by displacement from the epiblast rather than differentiation via trophectoderm.

The poor and aberrant colonisation behaviour of Dnmt3dKO cells prompted us to investigate in vitro differentiation competence. In response to mesendoderm induction, mutant cells up-regulated *T* and *Foxa2* but to a lower level than parental cells (Fig. S1C). In permissive conditions for neural induction (18), Dnmt3dKO cells showed only weak up-regulation of *Sox1* and *Pax6* (Fig. S1D) but displayed substantial and sustained up-regulation of *Ascl2, Hand1* and *Tpbpa*, transcription factor genes associated with the trophoblast lineage (Fig. S1C, S1D). To assess whether Dnmt3dKO cells adopt trophoblast identity, we applied two culture conditions for trophoblast cells (19, 20). Unlike parental cells, Dnmt3dKO cells showed no or low up-regulation of formative pluripotency factors Otx2 and Oct6, but instead gained expression of trophoblast markers (Fig. 1D, S1E).

We also subjected single *Dnmt* KO ES cells (15) to lineage induction (Fig. S1F). Expression of germ layer markers was reduced in both mutants and in neural conditions trophoblast genes were up-regulated (Fig. S1F). The phenotype was more marked in *Dnmt3b* KO cells, with higher trophoblast gene induction and lower neural and mesendodermal gene activation. We introduced expression constructs for *Dnmt3a* and *Dnmt3b* into Dnmt3dKO cells (Fig. S1G). Rescued cells displayed normal differentiation with suppression of trophoblast genes (Fig. S1H)

Culture in activin, FGF2 and KSR (AFK) induces ES cell conversion into post-implantation formative epiblast-like cells (EpiLCs) (21). Dnmt3dKO cells showed delayed morphological change on day 1 but by day 2 appeared similar to parental cells with a comparable increase in cell number (Fig S1I, S1J). The Rex1::GFPd2 (RGd2) reporter allows reliable monitoring of ES cell exit from naïve pluripotency (9). Dnmt3dKO cells showed delayed down-regulation of the reporter in AFK (Fig. S1K), consistent with findings in N2B27 (15). We sorted the GFP low (GFP^lo^) fraction that has exited naïve pluripotency and saw that trophoblast genes *Ascl2* and *Tfap2c* were mis-expressed in mutant cells (Fig. 1E). Levels increased further upon continued culture in N2B27 (Fig. 1E) while neural markers, *Sox1* and *Pax6*, were very lowly expressed (Fig. S1L). GFP^lo^ cells transferred into activin and CH gained only modest upregulation of *T* and FoxA2, though did not express most trophoblast markers.

After ongoing culture of dKO cells in N2B27, we observed additional trophoblast markers. Strikingly, however, the sequence of marker appearance differed from the in vivo developmental programme. *Ascl2* and *Hand1*, evident after 48h in AFK, are characteristic of post-implantation trophoblast, whereas *Cdx2* and *Elf5*, associated with primary trophectoderm, showed appreciable expression only at later stages (Fig. 1E). These results also differ from *Dnmt1* mutants where up-regulation of *Elf5* is thought to drive trophoblast transdifferentiation (22).

Finally, we transferred Dnmt3dKO cells from AFK into trophoblast medium (20). In contrast to parental cells, mutant cells flattened and some became very large with prominent nuclei (Fig. 1F). Propidium iodide staining showed a fraction with greater than 4N DNA content, consistent with polyploid trophoblast giant cell formation (Fig. 1G).

### Transcriptome analyses of ES cell differentiation trajectory in the absence of Dnmt3a/b

We examined the initial mis-regulation of gene expression by single cell RT-qPCR. We used *Nanog* (naive), *Otx2* (formative), and *Ascl2* (trophoblast) as representative markers (Fig. 2A, S2A). The majority of parental cells down-regulated *Nanog* and gained *Otx2* after 48h in AFK. Dnmt3dKO cells similarly gained Otx2 but generally retained higher Nanog levels. Most strikingly, *Ascl2* was upregulated in more than half of the Dnmt3dKO cells and many of these were also positive for both Nanog and Otx2. Unexpectedly, we also detected a fraction of triple positive cells among parental cells (Fig. 2A, S2A).

**Figure 2.**
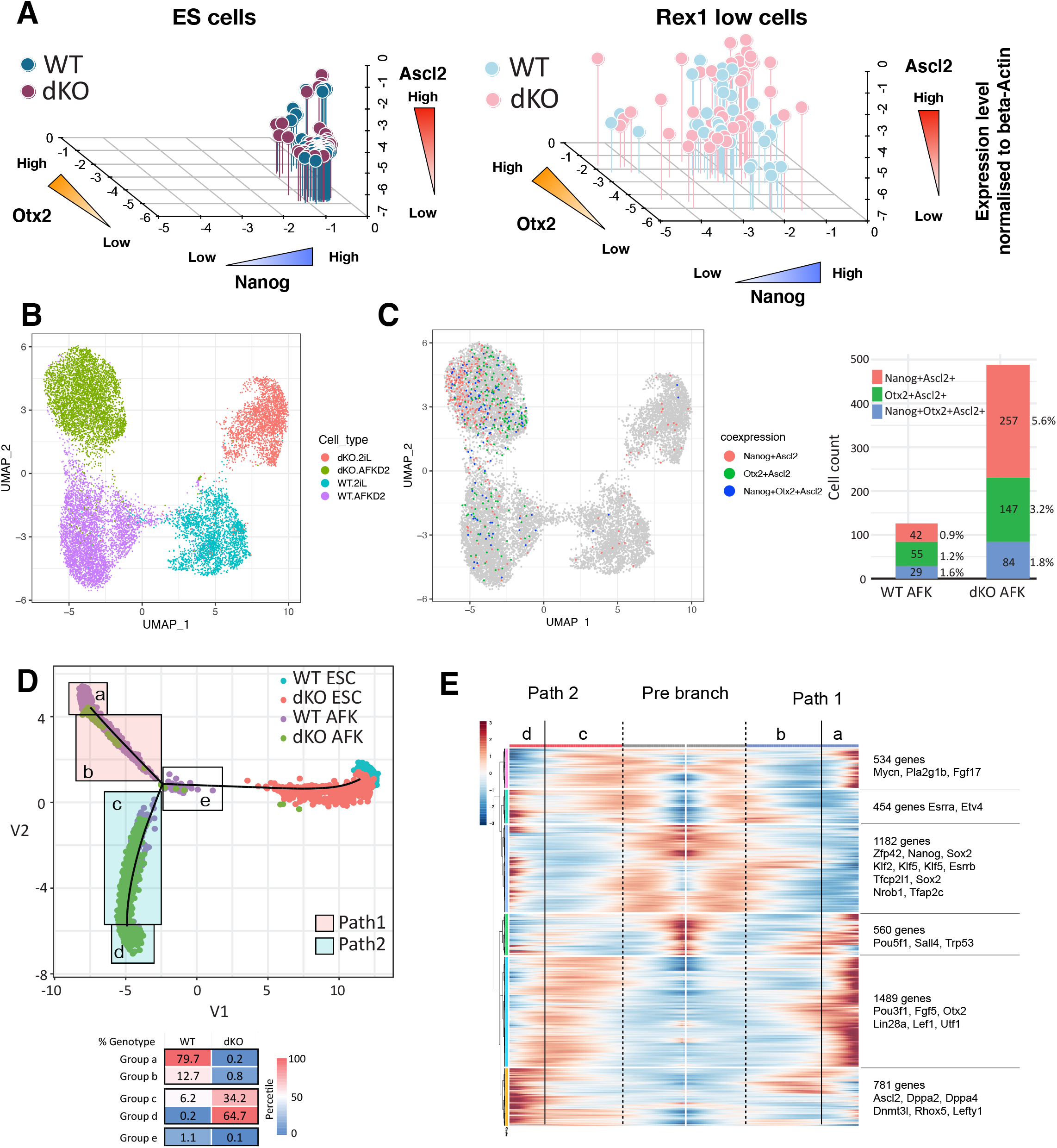
*Dnmt3a/b*-deficient ES cells adopt a deviant fate trajectory to post-implantation trophoblast. (A) Single cell qRT-PCR analysis of ES cells and sorted Rex1::GFP low AFK cells. Expression levels of Nanog (X axis), Otx2 (Y axis) and Ascl2 (Z axis) were normalized to beta-Actin. (B) UMAP of parental and dKO ES cells and AFK 48h cells. (C) UMAP colored to show cells co-expressing Nanog^+^Ascl2^+^ (red), Otx2^+^Ascl2^+^ (green) or Nanog^+^Otx2^+^Ascl2^+^ (blue) (left) with enumeration (right). (D) Pseudo-time ordering. Proportions of each genotype of AFK 48h cells in areas a-e are shown below. (E) Gene expression heatmap of 5,000 differentially expressed genes in pseudotime.

We extended this analysis using the 10x Genomics platform for single cell RNA-sequencing (scRNA-seq). UMAP analysis clustered cells by culture condition in the first dimension and genotype in the second dimension (Fig 2B). We inspected markers for pluripotency states, germ layers and trophoblast (Fig. S2B). In ES cells expression was not significantly altered between parental and mutant. However, in mutant 48h AFK cells we observed persistent expression of multiple naïve genes, lower up-regulation of formative genes, and ectopic expression of several trophoblast genes, though not Elf5 or Cdx2 which were detected in only 0.1% and 0.7% of cells respectively. We examined *Nanog, Otx2* and *Ascl2* using a UMI count above zero to classify expression. *Nanog*^+^*Ascl2*^+^, *Otx2*^+^*Ascl2*^+^ and *Nanog*^+^*Otx2*^+^*Ascl2*^+^ cells were present in both parental and Dnmt3dKO cells at 48hr (Fig. 2C). However, the combined proportion of dual and triple positive cells involving Ascl2 was three times higher in the mutants, consistent with single cell RT-qPCR.

Pseudo-time analysis using Monocle 2 (23) indicated a branchpoint in differentiation trajectory (Fig. 2D). We arbitrarily partitioned cells at and after the branchpoint into 5 groups (a-e). Parental cells were predominantly distributed along path 1 whereas mutant cells were almost exclusively located on path 2. Notably, however, 6.4 % of parental cells initiated path 2, although very few reached the endpoint. Differentially expressed genes in parental cells featured formative markers on path 1 and trophoblast genes on path 2 (Fig. S2C). Differential expression analysis without considering genotypes substantiated these alternative fates (Fig. 2E).

We investigated relatedness between path 2 and in vivo trophoblast. From published data (24) we identified genes upregulated in E3.5 trophectoderm or E6.5 trophoblast relative to inner cell mass and post-implantation epiblast respectively. Correlation was low for E3.5 trophectoderm-enriched genes with either pathway. In contrast, many E6.5 trophoblast-enriched genes were upregulated on path 2 (Fig. S2D).

### Chromatin accessibility and Ascl2 misexpression in the absence of de novo DNA methylation

Chromatin is remodelled during formative transition (25). We used ATAC-seq (26) to survey chromatin accessibility in the absence of de novo DNA methylation. We analysed ES cells, 48h GFP^Hi^ transitional cells, and GFP^Lo^ post-transition cells and identified loci that are more open at 48h in Dnmt3dKO cells than parental cells (Log2 fold change >0.7, P-value<0.05). We classified 4 groups according to opening in transitional (GFP^Hi^, Group I and III) or post-transition populations (GFP^Lo^, Group II and IV), and presence or absence of a CpG island (CGI), annotated by UCSC genome browser (Fig. 3A). The strongest peaks were associated with CGIs, which were lowly methylated at all stages (Fig. 3B and S3A, S3B). Non-CGI peaks were weaker and within regions that are methylated in parental cells, although a short stretch of reduced methylation was apparent at many Group IV peaks (Fig. 3B and S3A, S3B). As expected, CGI peaks are mainly associated with annotated promoters and non-CGI peaks with enhancers (Fig. S3C).

**Figure 3.**
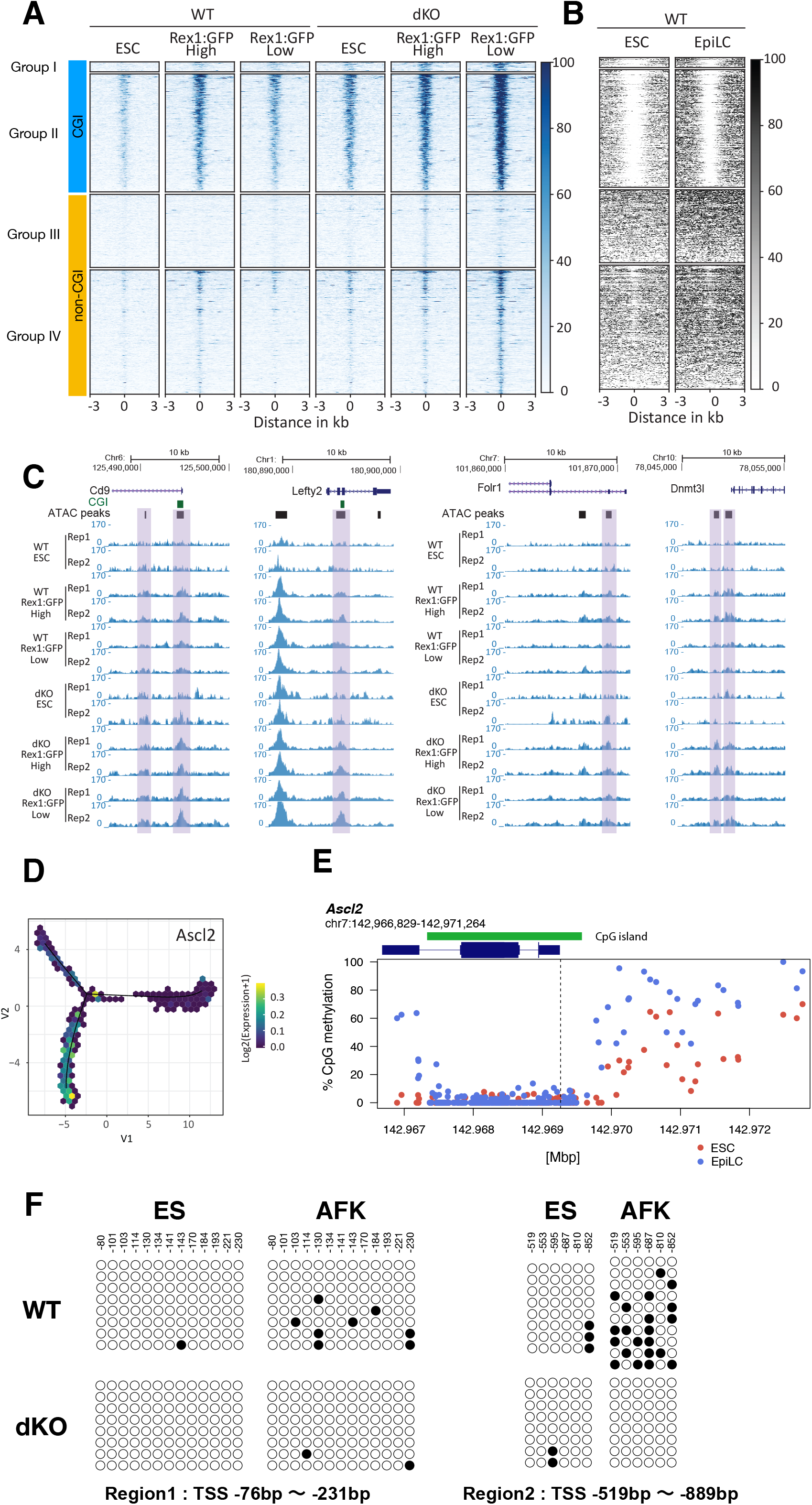
*Ascl2* locus opens during formative transition and remains open in *Dnmt3a/3b* mutants. (A) Heatmaps of ATAC-seq intensity distributions grouped according to increased accessibility in Dnmt3dKO GGP^Hi^ (I and III) or GFP^Lo^ (II and IV) presence or absence of CGI. (B) Heat map of CpG methylation across ATAC-seq peaks in ES cells and EpiLCs (10). (C) Genome browser examples of genes that are more open in Dnmt3dKO cells. (D)) Expression of *Ascl2* in pseudo-time. (E) *Ascl2* gene locus methylation pattern during ES to EpiLC transition (data from (10)). Dashed line marks the TSS. (F) Bisulfite Sanger sequence analysis of *Ascl2* TSS upstream sequence. Region 1 is within CGI and region 2 is within CGI shore. Filled circles represent methylated cytosine and open circles represents unmethylated cytosine. At least 8 clones each were sequenced.

The analysis revealed chromatin regions that opened transiently in parental cells but remained accessible in Dnmt3dKO cells (group II and IV, Fig 3A, S3B). Example genome browser tracks are shown in Fig. 3C. Genes associated with these peaks showed high correlation with differentially expressed genes in Dnmt3dKO GFP^Lo^ cells (Fig.S3D). We also found a positive correlation with the set of E6.5 trophoblast-enriched genes (Fig. S3E).

Thus, during exit from naïve pluripotency Dnmt3dKO ES cells fail to close down loci that normally open transiently during transition. These regions encompass promoters and proximal enhancers for a subset of E6.5 trophoblast genes that become mis-expressed.

Transcription factor motif enrichment analysis across ATAC peaks at CGI loci in Dnmt3dKO GFP^Lo^ cells identified, among others, Ascl1 (Fig. S3F). Ascl1 was not expressed in any of the samples studied, but the motif is shared with Ascl2, which, as noted above, is rapidly upregulated in transitioning mutant cells (Fig. 3E). The Ascl motif was present in 531 of 609 promoter regions (−2000 to +500bp around TSS) of E6.5 trophoblast enriched genes. In parental ES cells the *Ascl2* locus opened during formative transition (Fig. S3G) but in Dnmt3dKO cells *Ascl2* promoter accessibility increased further post-transition (Fig. S3F).

Inspection of published bisulfite sequencing data (10) revealed as expected that the *Ascl2* CGI is not methylated in ESCs or during naïve to EpiLC transition. However, the flanking CGI shore gained methylation during transition (Fig. 3F). We confirmed gain of CGI shore methylation in parental cells in AFK that did not occur in dKO cells (Fig. 3G). CGI shores are thought to contribute to regulation of CGI genes (27). The CGI shore sequence (1807 bp, within 2kb upstream of TSS) contains five Ascl2 motifs identified by JASPAR (28), consistent with auto-activation potential as reported in intestinal stem cells (29).

Previously it was found that ES cells deficient for Dnmt3a/3b gradually lost DNA methylation and after multiple passages could no longer form teratomas (14). That study was performed on ES cells cultured in serum, which have a hypermethylated genome compared to the early embryo. In 2i/LIF conditions used here the genome is hypomethylated, similar to the embryo (6-8). To examine the immediate effect of *Dnmt3a/3b* depletion we used selection for integration of gRNA and Cas9 expression vectors to rapidly isolate populations enriched for targeted cells. We transfected two ES cell lines and after three days cells were transferred to AFK for a further 48h. We detected up-regulation of *Ascl2*, albeit at a low level in these mixed populations (Fig. S3H). We carried out bisulfite sequencing of the *Ascl2* locus following *Dnmt3a/3b* deletion as above. We detected gain of methylation in the CGI shore in parental cells after 48h in AFK, that was reduced in the *Dnmt3a/3b* mutant population (Fig. S3I). These findings indicate a direct effect of *Dnmt3a/3b* depletion on both methylation and expression of *Ascl2*.

Imprinted silencing of the paternal allele of *Ascl2* in placenta does not involve methylation (30). Instead, the imprinted lncRN *Kcnq1ot1* suppresses transcription in *cis* (31). We inspected levels of *Kcnq1ot1* and found no change in Dnmt3dKO cells, indicating that loss of imprinting is not responsible for up-regulation of *Ascl2* (Fig S3J).

We then investigated whether expression of *Ascl2* is sufficient to impose trophoblast-like differentiation. For this we introduced an Ascl2-ER fusion construct into parental ES cells. Upon tamoxifen (Tx) treatment in serum and LIF, cells changed morphology within two days, becoming larger and flattened (Fig. 4A). *Oct4* was down-regulated and trophectoderm genes *Elf5, Cdx2* and *Tpbpa* were up-regulated (Fig. 4B). Thus, misexpression of *Ascl2* can initiate trophoblast-like differentiation.

**Figure 4.**
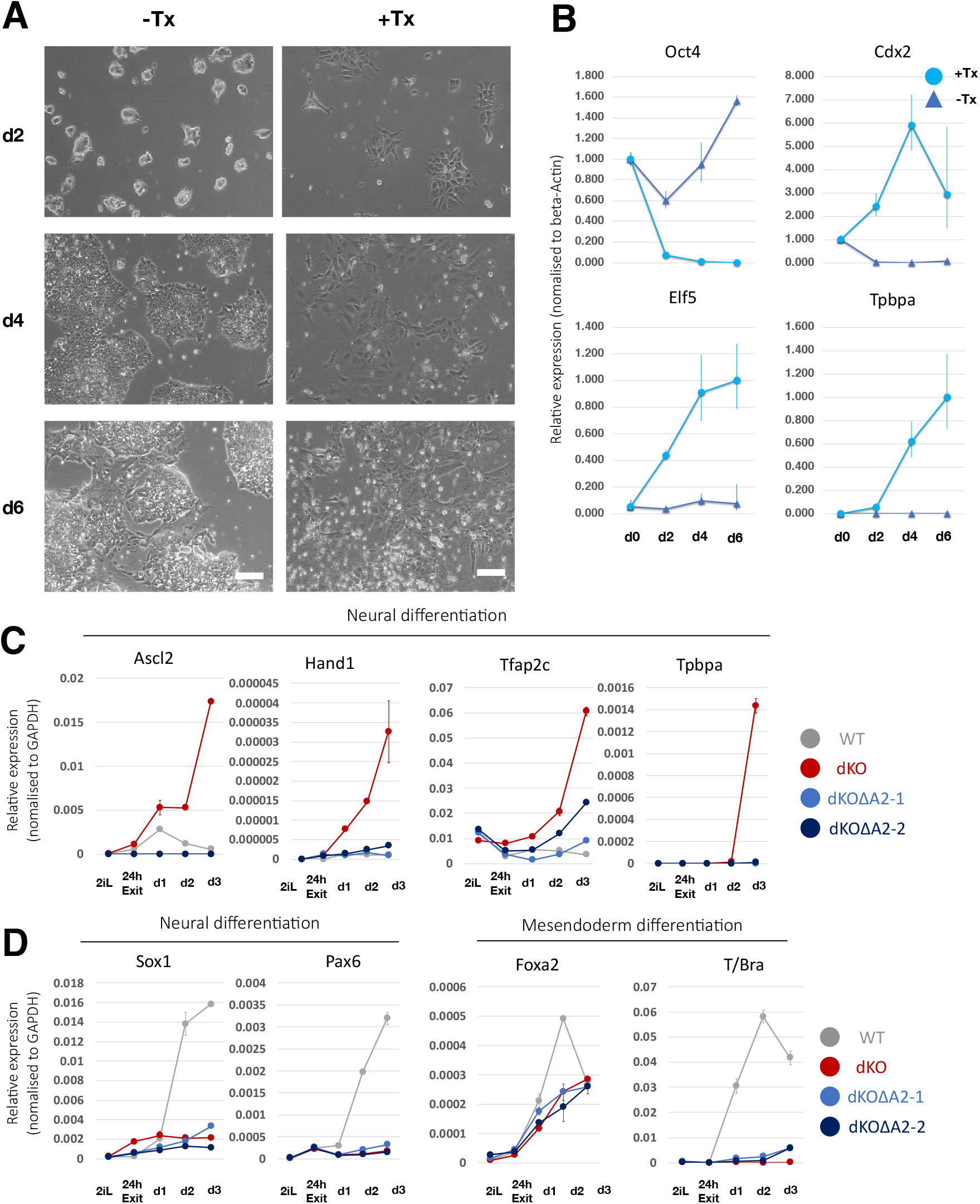
Ascl2 propels transdifferentiation to trophoblast-like cells. (A) Images of ES cells cultured in serum/LIF with or without tamoxifen (Tx) induced activation of Ascl2-ER. Scale bars, 100 µm. (B) qRT-PCR analysis of marker expression during Ascl2-ER induction. (C) qRT-PCR analysis of trophoblast marker expression in neural differentiation culture by two clonal lines of Dnmt3dKOΔA2 cells. (D) Analysis of somatic lineage marker expression by Dnmt3dKOΔA2 cells in neural and mesendoderm induction protocosl. Error bars represent S.D. from technical triplicates (B) or duplicates (C,D).

Finally, we tested whether activation of *Ascl2* was necessary for trophoblast-like differentiation of Dnmt3dKO cells. We deleted *Ascl2* to create Dnmt3dKOΔA2 cells (Fig. S4A) and saw that the mis-regulation of trophoblast genes was abolished (Fig. 4C). Furthermore, at 24h Dnmt3dKOΔA2 cells expressed *Otx2, Oct6* and *Fgf5* formative markers (Fig S4B). However, neural genes were not subsequently upregulated and mesendodermal differentiation remained inefficient (Fig. 4D), indicating later differentiation defects unrelated to Ascl2.

### Dnmt3a/b deficient cells are outcompeted by wildtype cells

We investigated whether deletion of *Ascl2* may restore contribution to chimaeric epiblasts. We saw substantial colonisation by Dnmt3dKOΔA2 cells in 5 out of 10 epiblasts at E6.5 (Fig. 5A, S5A). Moreover, we observed no donor cells in extraembryonic regions. At E7.5 contributions were reduced and appeared biased to the posterior region (Fig. 5B, S5B). Immunostaining for Stella and Tfap2c showed no enrichment for primordial germ cells (Fig S5C, D). By E9.5, we observed only sparse contributions in 4 of 11 embryos (Fig. 5C, S5E).

**Figure 5.**
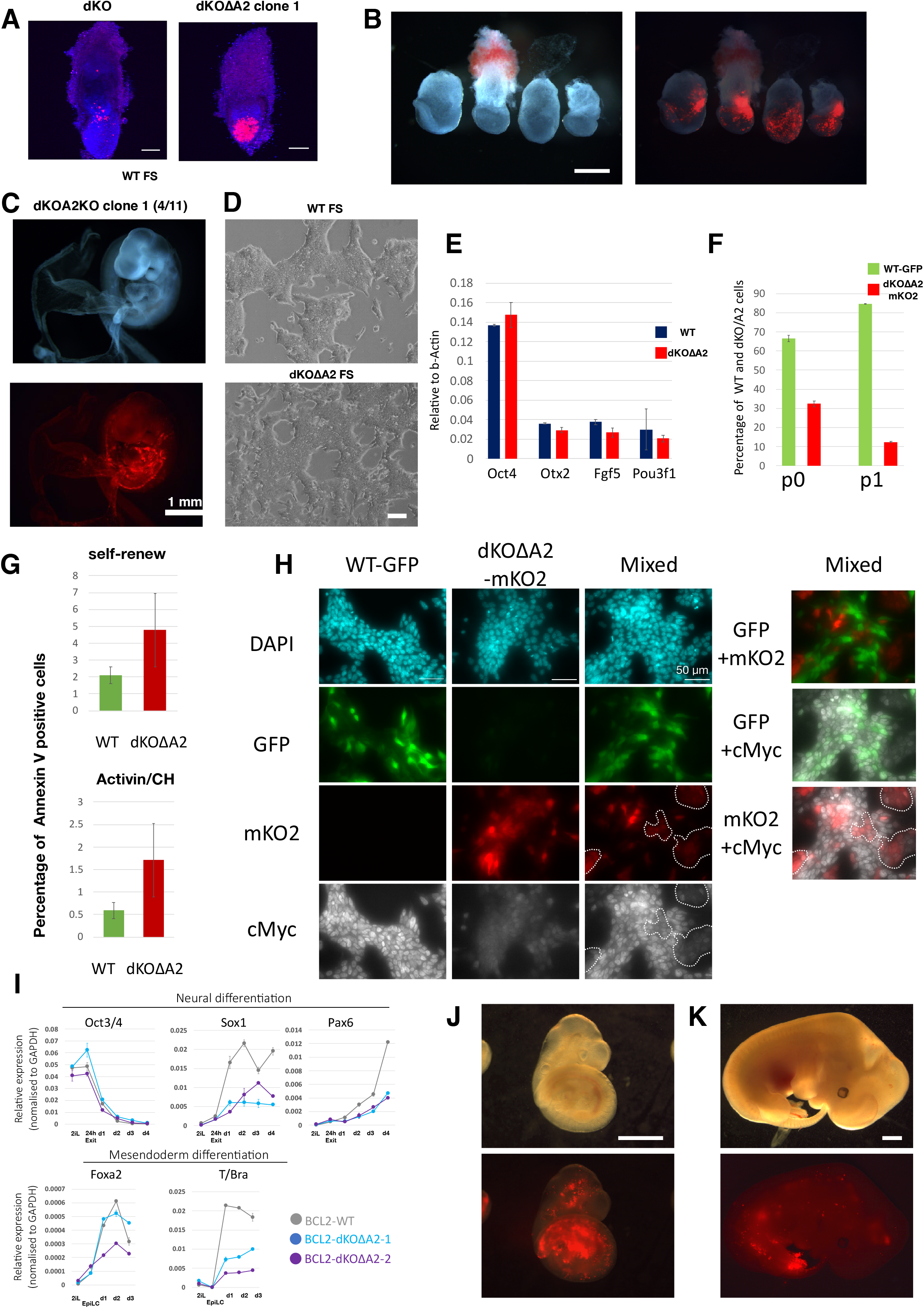
*Dnmt3a/b*-deficient cells are out-competed by wildtype cells and do not persist in chimaeras. (A) Maximum projection confocal images of chimeric contributions to E6.5 embryos. Red, mKO2; blue, Eomes immunostaining. Scale bar, 100μm. (B) Dnmt3dKOΔA2 ES cell chimaeras at E7.5. 25% opacity bright field image overlaid with fluorescent image. Scale bar, 250µm. (C) Dnmt3dKOΔA2 ES cell chimaeras at E9.5. Scale bar, 1mm. (D) FS cells established from parental and Dnmt3dKOΔA2 ES cells. Scale bar, 100µm. (E) RT-qPCR analysis of FS cell markers. Error bar represents S.D. from technical duplicates. (F) Percentage of GFP-labelled WT FS cells and mKO2-labelled dKO-A2 FS cells in the mixed co-culture. p0 is two days after plating. Error bar represents S.D from two cultures. (G) Annexin V positive cells quantified by flow cytometry after 24h in indicated co-culture. Error bars represents SD from 6 experiments. P<0.05. (H) Immunofluorescence images of cMyc parental FS cells (GFP), Dnmt3dKOΔA2 FS cells (mKO2) and co-culture for 24h. Dashed regions highlight mKO2 positive cells in co-culture. (I) RT-qPCR analysis of marker expression during neural and mesendoderm differentiation of BCL2 transfected ES cells. Error bar represents S.D. from technical duplicates. (J) BCL2-Dnmt3dKOΔA2 ES cell chimaeras at E9.5. 20% opacity bright field image overlaid on fluorescent image. Scale bar, 1mm. (K) BCL2-Dnmt3dKOΔA2 ES cell chimeras at E12.5. Scale bar, 1 mm.

Progressive dilution of mutant cells in chimaeras suggested a competitive disadvantage with wild type host epiblast cells (32). To examine this possibility we used formative pluripotent stem (FS) cells related to E6.0 epiblast (33). We were unable to establish FS cells from Dnmt3dKO cells, likely due to disruptive effects of *Ascl2* misexpression. However, Dnmt3dKOΔA2 ES cells could be converted into stable FS cell lines that expressed the core pluripotency factor Oct4 together with formative markers (Fig. 5D, E). We co-cultured equal numbers of GFP expressing wild type and mKO2 expressing Dnmt3dKOΔA2 FS cells. Proportions of each genotype were quantified by flow cytometry. We observed reduction in the fraction of mutant cells by passage 0 that was further increased by passage 1 (Fig. 5F). In co-cultures Annexin V staining was higher for Dnmt3dKOΔA2 FS cells during self-renewal and mesendoderm induction (p<0.05) though not neural induction (p>0.05) (Fig. 5G, S5G).

Relative level of cMyc is a key determinant in cell competition (34-36). By immunostaining we saw that Dnmt3dKOΔA2 FS cells have lower levels of cMyc protein than wild type cells (Fig. 5H). Immunoblotting confirmed that cMyc expression was reduced in mutant cells (Fig. S5F). These observations suggest that cMyc-based cell competition may cause elimination of Dnmt3dKO cells mixed with wild type cells (35) and that inactivation of *Ascl2* avoids only the first round of competition. The mechanism that reduces cMyc is unclear, but we surmise is secondary to other perturbations in the absence of de novo methylation.

We introduced *BCL2* into Dnmt3dKOΔA2 ES cells. Under in vitro differentiation conditions *BCL2* transfectants showed up-regulation of *Sox1, Pax6, T and Foxa2*, although still below the levels observed for parental transfectants (Fig. 5I). After blastocyst injection we recovered 5 chimaeras among 7 embryos at E9.5. Contributions were variable, but substantial in 3 of the 5 (Fig. 5J). We also collected embryos at E12.5 and detected mKO2 fluorescence in 2 out of 4 specimens, with donor-derived cells in neural tissues and mesenchyme (Fig. 5K, S5H). These contributions were noticeably lower than at E9.5, however, suggesting ongoing loss of *Dnmt3a/b*-deficient cells even in the presence of a strong survival factor.

## DISCUSSION

Our findings indicate that de novo DNA methylation safeguards the formative pluripotency transition to somatic lineage competence. In the absence of Dnmt3a and Dnmt3b, cells exiting naïve pluripotency are liable to adopt a deviant fate trajectory and develop features of post-implantation trophoblast. Notably, a small fraction of parental ES cells initiates the trophoblast transdifferentiation trajectory but they do not continue on this path because de novo DNA methylation prevents full activation of the trophoblast programme. Our study identifies Ascl2 as a trophoblast determination gene that is potentially directly regulated by de novo DNA methylation. Misexpression of Ascl2 can provoke ES cell transdifferentiation. Conversely, removal of *Ascl2* eliminates the aberrant fate trajectory in *Dnmt3a/3b* mutant cells and restores pluripotency progression. *Dnmt3a/3b*-deficient cells remain compromised in later differentiation, however, and are unable to compete with wildtype cells in the chimaera context.

Deletion of *Dnmt3a/3b* in ES cells generated in serum and LIF culture was previously shown to result in progressive erosion of methylation at repetitive sequences and, after long-term culture, a failure to produce teratomas (14). Effects of continuous culture on transcription or lineage commitment were not characterised. ES cells in serum are subject to heterogeneous and dynamic hypermethylation (37) whereas in 2i/LIF the genome is hypomethylated, similar to the inner cell mass (7, 38). The requirement for de novo methylation may be more acutely apparent in ground state ES cells in 2i/LIF due to the lower basal level of methylation. Crucially, the phenotype of *Ascl2* misregulation and trophoblast-like differentiation is specifically associated with absence of the de novo Dnmts and is eliminated by their restoration. The CGI shore adjacent to the *Ascl2* promoter is subject to de novo DNA methylation during formative transition, suggestive of a potential direct silencing effect. Alternatively, an upstream activator of *Ascl2* may be silenced by de novo methylation. Regardless of the mechanism, inspection of transcriptome data from *Dnmt3a/3b* deficient embryos at E8.5 (39) showed low but significant (P<0.01) up-regulation of *Ascl2*. Nonetheless, ectopic trophoblast differentiation is not the major cause of lethality in dKO embryos. Heightened susceptibility of ES cells to this fate alteration may reflect adaptation to the in vitro environment or the absence of constraints operative in the embryo. It is important to note that deregulation of *Ascl2* does not instruct normal trophoblast lineage development but triggers a deviant differentiation process and generation of an aberrant cell phenotype. Our findings serve as a cautionary note for interpretation of lineage potential and differentiation from pluripotent stem cells, in particular claims of expanded potency after exposure to epigenome modifying agents or other perturbations.

Overall, our results demonstrate that in the absence of de novo DNMTs the ability to execute a cell state transition is compromised due to misexpression of a gene with fate switching potency. The example of *Ascl2* illustrates how genome-wide methylation may have been co-opted to constrain expression of a pivotal gene that is transiently accessible during chromatin reconfiguration. This scenario may replay at other critical loci that open incidentally in the course of cell transitions. Indeed, transcriptome analysis of *Dnmt3a/3b* dKO embryos has highlighted deregulated expression of lineage-specific genes at E8.5 (39). The multiple abnormalities in *Dnmt3a/3b* mutant embryos and the inability of mutant ES cells to persist in any lineage (38) might be explained by a recurrent requirement for DNA methylation at critical loci to safeguard transcriptome trajectories. In this context it is of interest that mutations in *DNMT3a* and *DNMT3b* are associated with tumourigenesis in various tissues (40, 41), possibly relating to corruption of cellular transitions.

## MATERIALS AND METHODS

### Cell Culture

ES cells were maintained in 2iL medium on gelatin coated plates as described (18). 2iL medium consists of 10ng/ml mouse LIF, 1µM of Mek inhibitor PD0325901 and 3µM of GSK3 inhibitor CHIR99021 in N2B27 basal medium. EpiLCs were induced by plating 2×10^5^ ES cells in 20ng/ml activin A, 12.5ng/ml bFgf and 1% knockout serum replacement (KSR), in N2B27 medium on fibronectin coated 6-well plate. GFP high or low fractions were collected using MoFlo (Beckman Coulter) or FACS Fusion (BD) instruments. Neural differentiation was induced by plating 1×10^5^ cells in N2B27 basal medium on laminin coated 6-well plate. Mesendoderm induction was performed by plating 1×10^5^ cells in 10ng/ml Activin A and 3μM CHIR99021 in N2B27 medium on fibronectin coated 6-well plate. Trophoblast culture media were formulated as described previously (19, 20). For chimaera studies, cells were stably transfected with pPBCAG-mKO2-IP plasmid by TransIT LT1 (Mirus) with pCAG-PBase plasmid and selected with 1 µg/ml puromycin. Human *BCL2* was introduced by pT2PyCAG-hBCL2-IH by TransIT LT1 with pCAG-T2ase plasmid and selection with 100µg/ml of hygromycin B. To establish tamoxifen inducible *Ascl2* expression lines, pPBCAG-Ascl2ER-IN plasmid was co-transfected with pCAG-PBase plasmid into E14Tg2a ES cells and clones picked after selection with 250µg/ml G418. DnmtdKOΔA2 FS cells were established by conversion of ES cells and maintained in 3ng/ml of activin A, 2µM XAV939 and 1µM BMS493 in N2B27 medium on fibronectin coated plates as described (33). Alexa Fluor 647 conjugated Annexin V (Biolegend, Cat#640911) was used for apoptotic cell detection by FACS Fortessa (BD).

### Gene targeting

*Dnmt3a* or *Dnmt3b* single KO and double KO cells were established previously using CRISPR/Cas9 and gRNAs designed to excise sequences encoding the catalytic domain (15). Knockout cells were used within 15 passages and compared with wild type cells of similar passage. To assess the effect of gene knockout acutely, gRNAs were cloned into a piggyBac vector with puromycin resistance cassette, pCML32 (33), and co-transfected with pPB-Cas9-IN and pCAG-PBase using TransIT-LT1 (Mirus) followed by selection with 1µg/ml Puromycin and 300µg/ml G418. To rescue Dnmt KO cells, pPB-CAG-Dnmt3a-IP and pPBCAG-Dnmt3b-IZ were co-transfected with pCAG-PBase usingTransIT-LT1 with selection in 1 µg/ml Puromycin and 40 µg/ml Zeocin. *Ascl2* gRNAs were designed to excise Exon2, which encodes the full-length protein. Clones were screened by genomic PCR. gRNAs and primers are listed in Table S1.

### Chimaeric embryo experiments

Reporter expressing ES cells were dissociated and 10-15 cells injected into each blastocyst. Injected embryos were transferred to the uteri of pseudo-pregnant females or cultured in vitro for 24h in M2 medium (Sigma Aldrich) in 7% CO_2_ at 37°C.

### Immunostaining

Embryos and culture cells were fixed with 4% PFA at RT (for 15 minutes for cells and upto E7.5 stage embryos, 60 minutes for E9.5 and E12.5). Whole E6.5 embryos were stained with rabbit anti-Eomes antibody (Abcam, Cat#ab23345). Blastocyst were stained with rabbit anti-GFP (Invitrogen, Cat#A-11122) and rat anti-Sox2 (eBioscience, Cat#14-9811-82). After the fixation, E9.5 and E12.5 embryos were incubated with 20 % sucrose/PBS overnight at 4°C, then embedded in OCT compound. Sections were stained with rat anti-Sox2 (eBiosciencce) and FITC-conjugated mouse anti-cTnT antibodies (Abcam, Cat#ab105439) (E9.5 only). Nuclei were stained with DAPI. Embryos and sections were imaged by Leica SP5 or Zeiss LSM880 confocal microscope. E7.5 PGC were stained with anti-Tfap2c (Santa Cruz, Cat# sc-8977), anti-Stella (Abcam, Cat#ab19878) and rat anti-Sox2 (eBioscience, Cat#14-9811-82). FS cells were stained with rabbit anti-cMyc antibody (Abcam, Cat#ab30272).

### Western Blot

Cells were lysed with RIPA buffer in the presence of Protease/Phosphatase inhibitor cocktail (Invitrogen). Lysed cells were rotated for 20 minutes and sonicated in Bioruptor (Diagenode). Cell lysates were cleared by centrifugation, and the supernatant was recovered. Protein concentrations were measured by the BCA method (Pierce). 15 µg of protein was loaded in each well. Blots were blocked with 5% BSA/TBS 0.1 % Triton-X for 1 hour at RT and incubated overnight with primary antibodies at 4°C. Secondary antibodies were incubated for 1 hour at RT and signals were detected with ECL Select (GE Healthcare) and Odyssey Fc (Li-Cor). NaOH (0.2N) was used for stripping. Anti-cMyc (Abcam, Cat#ab30272) and anti-αTubulin (Thermo Fisher Scientific, A-11126) were used.

### qRT-PCR

Total RNA was isolated with Relila RNA miniprep systems (Promega) and cDNA was synthesized using GoScript Reverse Transcriptase system (Promega). qRT-PCR was performed with Taqman Gene Expression (Thermo Fisher Scientific) or Fast SYBR Green Master Mix (Thermo Fisher Scientific). Primers are listed in Table S1.

### Bisulfite Sanger sequencing analysis

Genomic DNA was prepared with DNeasy Blood and Tissue Kit (Qiagen). Purified genomic DNA was treated with Imprint DNA modification kit to perform bisulfite conversion. Ascl2 promoter regions are amplified by Nested PCR with Touchdown protocol with LongAmp DNA Taq polymerase (New England Biolabs). The PCR for both rounds were performed as follows; denaturing at 94°C 30sec, 10 cycles of gradient PCR, 94°C for 15 sec, 65°C (annealing temperature reduced 1°C per cycle) for 15 sec, 65°C for 30 sec and 35 cycle of 94°C for 15 sec, 56°C for 15 sec and 65°C for 30 sec. 2µl of first PCR product was used for the nested PCR. PCR product were purified from agarose gel and cloned into TOPO cloning vector (Thermo Fisher Scientific). The sequence results were analysed with QUMA (42).The primers are listed in Table S1.

### Single cell qPCR

Custom Taqman probes were ordered for Nanog, Otx2, Ascl2 and Actin-beta (Thermo Fisher Scientific). Single cells were collected directly into 96-well plate by flow cytometry (MoFlo, Beckman Coulter) and reverse transcription was performed as described previously (43). Cells which had Ct-value of Actin-beta >14 and no amplification were excluded from the analysis.

### Single cell RNA-seq

ES cells and 48 hours AFK cells were dissociated with accutase (Biolegend) and live cells were collected by flow cytometry (MoFlo, Beckman Coulter). Single cell cDNA libraries were constructed using 10x Genomics technology and sequenced by the Genomics Core Facility, Cancer Research UK Cambridge Institute.

### ATAC-seq

50,000 ES cells and AFK cells at 48h were collected by flow cytometry (MoFlo, Beckman Coulter). Cells washed with ice-cold PBS once then lysed in the buffer (10 mM Tris-HCl, pH 7.4, 10 mM NaCl, 3 mM MgCl2 0.1% IGEPAL). The nuclear pellets were collected and Tn5 tagmentation and library construction were performed with the Illumina Nextera kit (FC-121-1030). DNA was purified with AMPure XP beads (Beckman Coulter).

## DATA ANALYSIS

### Single Cell RNAseq analysis

Preliminary sequencing results (bcl files) were converted to fastq files with CellRanger (version 3.0) following the standard 10x Genomics protocol. Barcodes and unique molecular identifier (UMI) ends were trimmed to 26 bp, and mRNA ends to 98 bp. Reads were then aligned to the mouse reference genome (mm10) and gene counts were obtained using the GRCm38.92 annotation in CellRanger. We used Seurat (version 3.1.5) (44) to further process the single-cell RNA-seq data. Cells that have between 200 and 5500 unique gene counts, and with less than 5% mitochondrial gene content were filtered. We analysed 2,724 parental and 2,459 Dnmt3dKO ESCs with 4,776 parental and 4,563 Dnmt3dKO 48hr cells. Log-normalization using a scale factor of 10,000 was performed, and the data were scaled to produce standardized expression values for each gene across all cells (z-score transformation), while also regressing out unwanted variation in the percent of mitochondrial gene content. We determined the top 2,000 most highly variable genes, and performed principal component analysis. Visualisation was performed using the UMAP projection method on the first 18 principal components. Differential expression was performed using the Wilcoxon rank sum test, using a threshold of log2 fold change greater than 0.1 and p-value less than 0.05. The enrichment of TE (E3.5) and ExE (E6.5) genes was visualised using the AddModuleScore function of Seurat to show the average expression of sets of genes. Co-expression of Nanog, Ascl2, and Otx2 was assigned to cells having at least one unique read for each of the genes.

### Pseudotemporal analysis

Pseudotime trajectory was calculated using Monocle2 (version 2.14) (45-47) using the method “DDRTree” with the top 1000 differentially expressed genes between each of the clusters found from the Seurat clustering. To find differences between cells at the end and middle of each branch, the trajectory was split into sections using an arbitrary pseudotime cut-off of 21.4 for path 1 and 22.25 for path 2 based on visualizing the trajectory. To find genes that are differentially expressed along pseudo-time the monocle function differentialGeneTest was used with default parameters.

### Bulk RNA-seq analysis

Genes enriched in trophoblast over epiblast at E6.5 or in trophectoderm over ICM at E3.5 were determined by differential expression analysis using publicly available data sets (GSE84236). This was done using tximport (48) to load the dataset into DESeq2 (48) for differential expression analysis using a threshold of log2 fold change > 2 and an adjusted p-value of less than 0.05.

### ATAC-seq analysis

Raw reads were pre-processed and quality-filtered using Trim Galore! (https://github.com/FelixKrueger/TrimGalore), and reads shorter than 15 nt were discarded. Processed reads were then aligned to the mouse reference genome (mm10) using bowtie with parameters “-m1 -v1 --best --strata -X 2000 --trim3 1”. Duplicates were removed using Picard tools. Read pairs larger than one nucleosome length (146 bp) were discarded, and an offset of 4 nts was introduced. Peaks were called with MACS2 and parameters “--nomodel --shift - 55 --extsize 110 --broad -g mm --broad-cutoff 0.1”. Then the R package Diffbind^8^(49) was used to calculate reads across the merged peaks and calculate differential peaks for each cell type utilising the edgeR method (50, 51). Peaks were also identified as being within a CpG island or not and assigned genes based on a window of 2kb around each peak. Scatterplots were plotted using the differential ATAC-seq fold changes from edgeR against the RNA-seq fold changes from Seurat. In each of the quadrants of the scatterplot Fisher’s exact test was used to calculate whether the differential expressed genes were overrepresented. Motif analysis was performed by using the findMotifsGenome tool in homer (version v4.10) (52) on peaks that overlapped with CpG islands in the KO using the WT peaks as background sites.

## Supporting information

Supplemental Table of Primers

## Data Availability

10xGenomics and ATAC-seq data are deposited in GEO: GSE158347.

## Acknowledgments

We thank Brian Hendrich and Jennifer Nichols for comments on the manuscript and Giuliano Stirparo for advice on informatics. We are grateful to Tim Lohoff and Jennifer Nichols for discussion of unpublished data. We thank Maike Paramor, Vicki Murray, Peter Humphreys, Darran Clements, Andrew Riddell and biofacility staff for technical support and the CSCI core bioinformatics team for data processing. Sequencing was performed by the CRUK Cambridge Institute Genomics Core Facility. Rosalind Drummond and James Clarke provided laboratory assistance. This research was funded by the Biotechnology and Biological Sciences Research Council (BB/P009867/1, BB/P021573/1) and the Medical Research Council (MR/P00072X/1). The Cambridge Stem Cell Institute receives core funding from Wellcome (203151/Z/16/Z) and the Medical Research Council (MC_PC_12009). AS is a Medical Research Council Professor (G1100526/1).

## Author Contributions

Conceptualisation, M.K., M.A.L., A.S.; Methods, M.K., M.A.L.; Formal Analysis, M.B., S.D.; Investigation M.K., M.A.L., W.M.; Writing M.K., M.A.L., A.S.; Supervision, A.S.

## Notes

### Competing Interest Statement

The authors have declared no competing interest.

### Summary of Updates

The revised version includes data showing gain of methylation of the Ascl2 CGI shore during formative transition that fails to occur in Dnmt3a/3b double mutant ES cells. We also provide evidence that Ascl2 is mis-regulated immediately following Dnmt3a/3b deletion. The abstract and discussion are updated and revised.

