## Supplemental figures and legends for "Disabling *de novo* DNA methylation in embryonic stem cells allows an illegitimate fate trajectory"

**SUPPLEMENTAL INFORMATION**

Fig. S1 (A) Maximum projection confocal microscopy images of BCL2 ES cell chimaeras in Fig 1C, immunostained for Eomes in blue. Scale bar, 100μm. (B) Representative images of blastocysts injected with hBCL2 parental or Dnmt3dKO ES cells and cultured for 24h. Scale bars, 50μm. (C) qRT-PCR analysis of neural lineage and trophoblast markers during neural induction. (D) qRT-PCR analysis of primitive streak and trophoblast markers during mesendoderm induction. (E) qRT-PCR analysis of trophoblast and epiblast markers after seven days in alternative trophoblast cell medium, FAXY (Ohinata and Tsukiyama 2014). (F) RT-qPCR analysis of neural and mesendodrm gene expression in *Dnmt3a* or *Dnmt3b* single mutant ES cells. (G) qRT-PCR assay of *Dnmt3a* and *Dnmt3b* expression levels in WT, dKO and Dnmt3a/b-transfected rescue dKO ES cells. (H) qRT-PCR analysis of somatic lineage and trophoblast marker expression in rescued dKO ES cells. (I) Morphology of WT and dKO cells after 24h and 48h in AFK. Scale bars, 100 μm. (J) Cell counts 48h after plating $2x10^5$ ES cells in AFK. Error bars represent SD from biological replicates (n=3). p>0.05. (K) Rex1::GFP flow cytometry profile during AFK culture for three days. (L) qRT-PCR analysis of 2iL ES cells, Rex1::GFP low 48h AFK cells, and cells differentiated for 3 days post-sorting. All qPCR data are normalized by beta-Actin and error bars represent S.D. from technical duplicates.

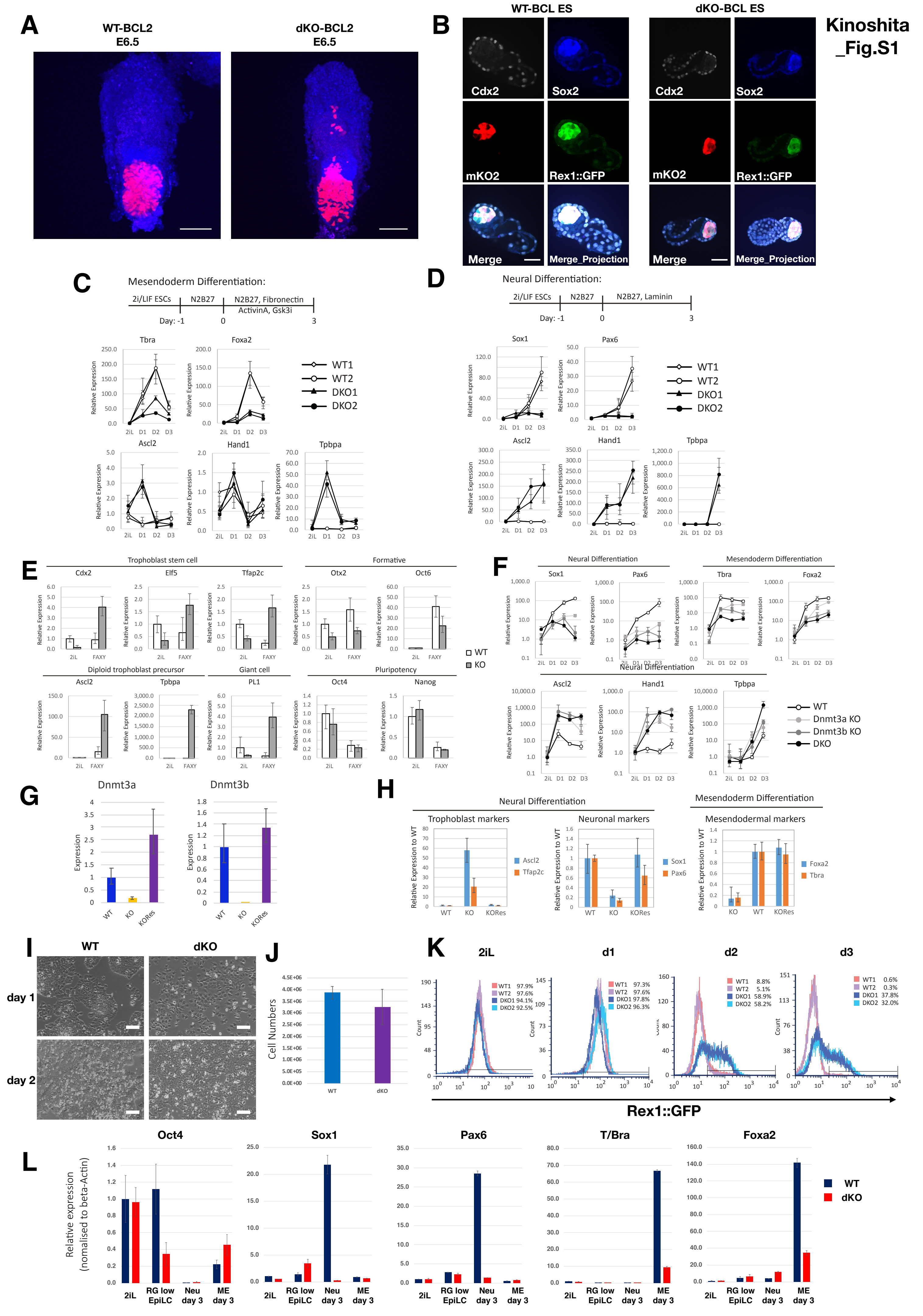

Kinoshita _Fig.S1
A
WT-BCL2 E6.5
dKO-BCL2 E6.5
B
WT-BCL ES
dKO-BCL ES
Cdx2
Sox2
mKO2
Rex1::GFP
Merge
Merge_Projection
C
Mesendoderm Differentiation:
2i/LIF ESCs
N2B27
N2B27, Fibronectin ActivinA, Gsk3i
Day: -1 0 3
Tbra
Foxa2
Ascl2
Hand1
Tpbpa
Relative Expression
2iL D1 D2 D3
WT1
WT2
DKO1
DKO2
D
Neural Differentiation:
2i/LIF ESCs
N2B27
N2B27, Laminin
Day: -1 0 3
Sox1
Pax6
Ascl2
Hand1
Tpbpa
E
Trophoblast stem cell
Cdx2
Elf5
Tfap2c
Formative
Otx2
Oct6
Diploid trophoblast precursor
Ascl2
Tpbpa
Giant cell
PL1
Pluripotency
Oct4
Nanog
2iL FAXY
WT
KO
F
Neural Differentiation
Sox1
Pax6
Mesendoderm Differentiation
Tbra
Foxa2
Neural Differentiation
Ascl2
Hand1
Tpbpa
WT
Dnmt3a KO
Dnmt3b KO
DKO
G
Dnmt3a
Dnmt3b
Expression
WT
KO
KORes
H
Neural Differentiation
Trophoblast markers
Ascl2
Tfap2c
Neuronal markers
Sox1
Pax6
Mesendoderm Differentiation
Mesendodermal markers
Foxa2
Tbra
Relative Expression to WT
I
WT
dKO
day 1
day 2
J
Cell Numbers
WT
dKO
K
2iL
d1
d2
d3
Count
WT1 97.9%
WT2 97.6%
DKO1 94.1%
DKO2 92.5%
WT1 97.3%
WT2 97.6%
DKO1 97.8%
DKO2 96.3%
WT1 8.8%
WT2 5.1%
DKO1 58.9%
DKO2 58.2%
WT1 0.6%
WT2 0.3%
DKO1 37.8%
DKO2 32.0%
Rex1::GFP
L
Oct4
Sox1
Pax6
T/Bra
Foxa2
Relative expression (nomalised to beta-Actin)
2iL
RG low EpiLC
Neu day 3
ME day 3
WT
dKO

Fig. S2 (A) Related to Fig. 2A. Single cell qRT-PCR expression levels of Nanog, Otx2 and Ascl2 in undifferentiated ES cells and Rex1-low AFK cells at 48h. (B) Marker gene expression from scRNA-seq data. Size of circles indicates percentage of cells expressing each gene and colour scale indicates expression level. (C) Heatmap of differentially expressed genes (Log2 fold change >0.1 and p-value<0.05) between parental cells in groups b and c from Figure 2d. (D) Expression pattern of E3.5 TE and E6.5 ExE enriched genes along the pseudo-time trajectories, Path 1 in blue and Path 2 in red. Solid lines are mean values. Enriched genes (expression Log2 fold change of 2 and adjusted p-value<0.05) were identified for trophectoderm vs ICM from E3.5 blastocysts and extraembryonic ectoderm vs epiblast from E6.5 embryos (data from Smith Z.D. et al 2017).

Kinoshita_Fig.S2

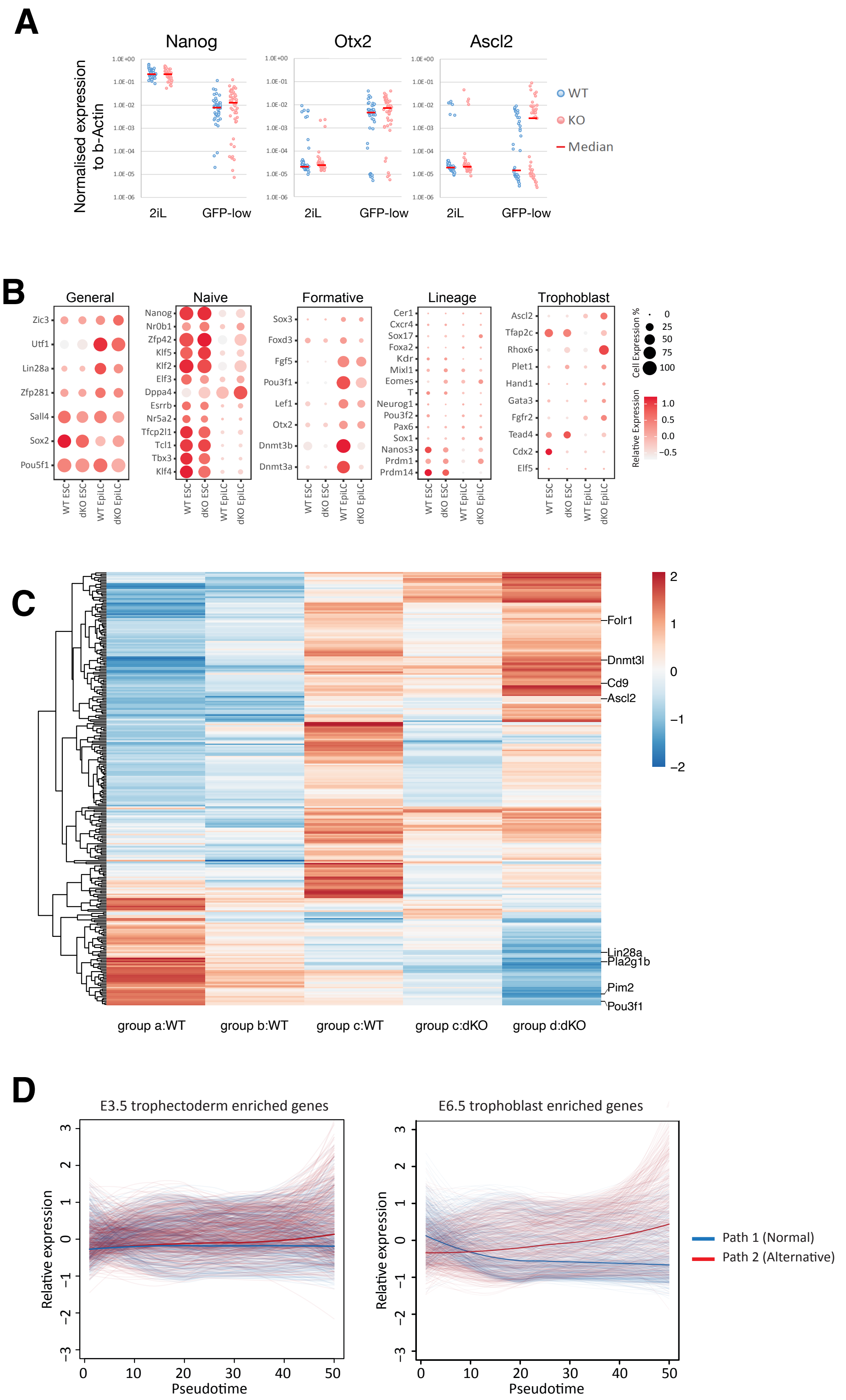

Fig. S3 (A) CpG DNA methylation pattern of identified peaks in WT ES AFK cells in Fig.3A. Averaged percentage of DNA methylation from each group is shown. (B) RPKM values from each group identified in Fig. 3A. (C) Association (within 2kb) of ATAC-seq peaks in groups I-IV promoters or enhancers. (D) Correlation plot for open chromatin regions in GFP low AFK cells with differentially expressed genes between WT and dKO AFK 48h cells. (E) Correlation plot for open chromatin regions with expression in AFK cells of E6.5 trophoblast-enriched genes associated with ATAC peaks. (F) De novo motif analysis of differential ATAC peaks. (G) ATAC-seq profile of *Ascl2* gene locus in WT and dKO cells in 2iL and AFK. (H) qRT-PCR analysis in ES cells and AFK day 2 cells immediately after Dnmt3a/3b depletion as depicted in the schematic. Results are from two independent ES cell lines. Error bars represent S.D. from technical duplicates. (I) CGI shore CpG methylation analysis after Dnmt3a/3b depletion as above in an independent experiment. Filled circles represent methylated cytosine and open circles unmethylated cytosine. qRT-PCR shows *Ascl2* gene expression level from the same samples. Error bars represent S.D. from technical duplicates. (I) UMAP from Fig 2B coloured to show *Kcnq1ot1* expression values.

**Kinoshita_Fig.S3**

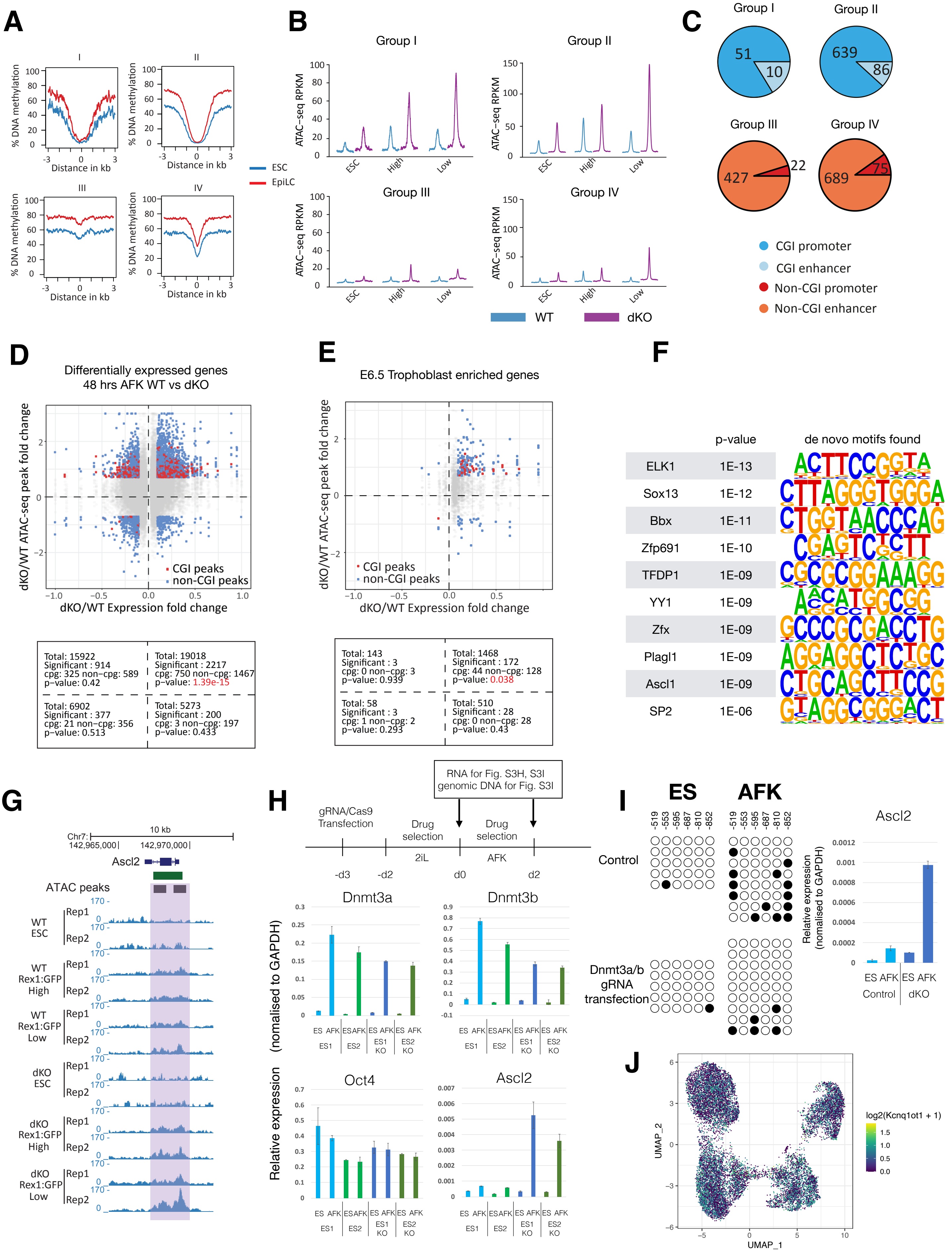

Fig.S4 (A) *Ascl2* KO genotyping by genomic PCR. Wildtype band is 2563 bp and Ascl2 KO band 412 bp. (B) RT-qPCR analysis of formative marker expression in indicated cells in 2iL and after 24h in N2B27. Error bars represent S.D. from technical duplicates.

Kinoshita_Fig.S4

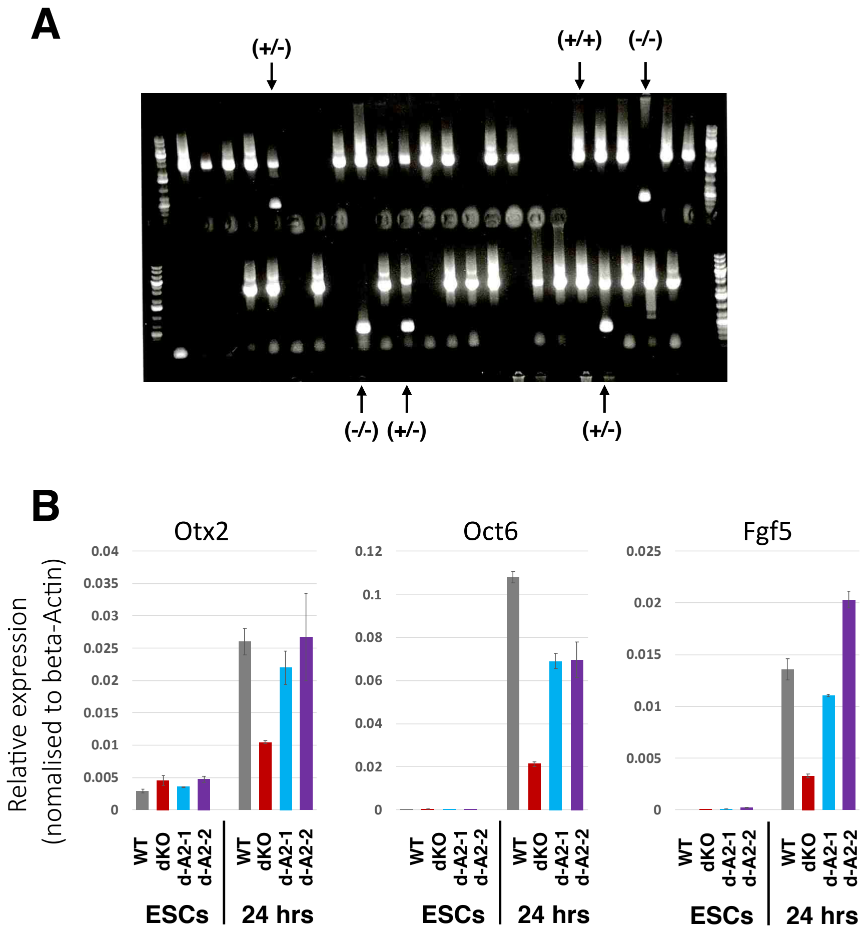

Fig. S5 (A) Related to Fig5A. Maximum projection images from z-stack confocal microscopy slices of E6.5 chimaeric embryos. Eomes staining is in blue and mKO2 reporter in red. Scale bar, 100μm. (B) E7.5 chimaeras from two mKO2 reporter Dnmt3dKOΔA2 ES cell lines. Scale bars, 1mm (left), 500μm (right). (C) E7.5 chimaeric embryo immunostained for Stella (green) and Sox2 (blue). Scale bar, 100 μm. (D) E7.5 chimaera immunostained for Tfap2c (green). (E) Section of E9.5 chimaeric embryo immunostained for Sox2 (blue) and cTnT (green). mKO2 (red) positive cells are present in neuroepithelium (white arrowhead), foregut (white arrow) and cardiac mesoderm (blue arrow). Scale bar, 250μm. (F) Western blot analysis of cMyc protein in WT and Dnmt3dKOΔA2 cells. Proteins were collected from independent WT or Dnmt3dKOΔA2 FS cell cultures (unsort) and by sorting GFP positive or mKO2 positive fractions from a mixed culture after 1 day (sorted). (G) Proportions of Annexin V positive cells of each genotype determined by flow cytometry after co-culture for 1 day in N2B27 medium. Error bars represents SD from 6 experiments. P>0.05. (H) Confocal microscope images of Sox2 immunostaining and mKO2 reporter expression in brain region of chimaera shown in Fig 5K. Scale bar, 100μm.

Kinoshita_Fig.S5

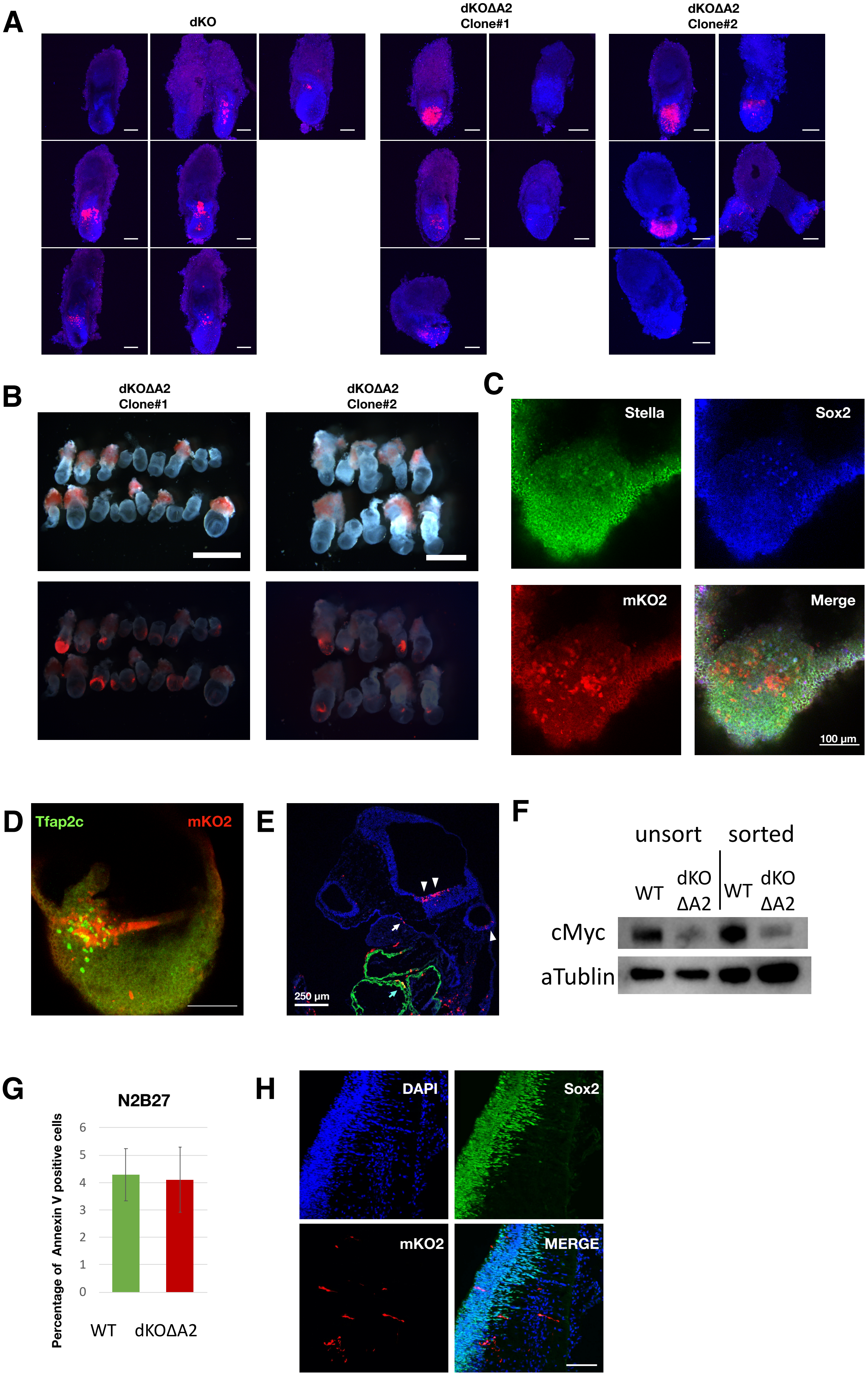
